# Alpha-synuclein overexpression induces sex-dependent alterations in striatal autophagic-lysosomal markers in a Parkinson’s disease mouse model

**DOI:** 10.64898/2026.11.27.769612

**Authors:** Raphaelle Bidgood, Maider Zubelzu, Erlantz Eguinoa, Isabel Jimenez-Ridruejo, Ane Murueta-Goyena, Jose Angel Ruiz-Ortega, Jose Vicente Lafuente, Nicolas P Mallet, Ainhoa Plaza-Zabala, Teresa Morera-Herreras

## Abstract

Parkinson’s disease (PD) is characterized by pathological alpha-synuclein (α-syn) accumulation and alterations in autophagy. While biological sex is increasingly recognized as a modifier of PD susceptibility and pathophysiology, its relationship with striatal autophagic-lysosomal alterations remains incompletely understood. In this study, we investigated sex-dependent changes in related molecular markers in a mouse model overexpressing α-syn. Male and female C57BL/6J mice received bilateral intranigral injections of adeno-associated viral vectors encoding mutant A53T α-syn or empty vectors. Dorsal striatal tissue from 29 animals was analyzed at 60 or 120 days post-injection. Confocal imaging of LC3B (microtubule-associated protein 1 light chain 3 beta; autophagosome marker) and LAMP1 (lysosomal-associated membrane protein 1; lysosomal marker) immunofluorescence quantified marker-positive areas, puncta densities, and colocalization. Associations with α-syn immunoreactivity, striatal dopaminergic innervation, and axonopathy severity were also examined. At 60 days, no vector-related differences in autophagic-lysosomal markers were detected. However, at 120 days, vector × sex interactions were identified for LC3-positive area, LAMP1-positive area, and LC3 puncta density. Post-hoc comparisons showed reductions in all three measures in α-syn males relative to male controls, whereas the corresponding comparisons in females were not significant. After adjustment for striatal α-syn immunoreactivity, α-syn males had lower LC3- and LAMP1-positive areas, LC3-LAMP1 colocalization area, and LAMP1 puncta density than α-syn females. Across α-syn mice, greater LAMP1 puncta density was associated with more advanced axonopathy after adjustment for sex and time. These findings identify sex-dependent alterations in striatal autophagic-lysosomal markers following A53T α-syn overexpression and suggest an association between lysosomal organization and axonopathy severity. They also underscore the importance of considering biological sex in studies of PD pathophysiology and when evaluating therapies targeting protein clearance pathways.

**IMPACT STATEMENT:** Alterations in the autophagy-lysosome system are implicated in alpha-synuclein (α-syn) accumulation and neurodegeneration in Parkinson’s disease, but their relationship with biological sex remains poorly understood. In an A53T α-syn mouse model, we identified sex-dependent changes in striatal autophagic and lysosomal markers. At 120 days post-injection, α-syn overexpression was associated with reduced LC3- and LAMP1-positive areas and lower LC3 puncta density in males, whereas these vector group changes were not detected in females. Several differences between α-syn males and females persisted after adjustment for striatal α-syn expression, suggesting that sex-dependent tissue responses to α-syn were not solely explained by α-syn levels. Greater LAMP1 puncta density was also associated with more advanced axonopathy after accounting for sex and time. These findings identify sex-related alterations in autophagic-lysosomal organization as a cellular correlate of differential vulnerability to α-syn pathology, and support considering biological sex when evaluating therapies targeting protein clearance.

## INTRODUCTION

Parkinson’s disease (PD) is a progressive neurodegenerative disorder characterized by degeneration of nigrostriatal dopaminergic neurons and accumulation of misfolded alpha-synuclein (α-syn)^1,2^. Although the mechanisms underlying neuronal loss remain unclear, disrupted cellular homeostasis is considered central to disease progression^3,4^. As post-mitotic cells, neurons depend heavily on intracellular degradation pathways to maintain proteostasis, making them particularly vulnerable to dysfunction in these systems^5-7^. Autophagy contributes to the clearance of α-syn and other damaged intracellular components by sequestering cargo in autophagosomes that fuse with lysosomes for degradation^8^. Impairments in autophagosome formation, lysosomal function, and substrate degradation have been reported in experimental and *postmortem* PD studies^6, 9, 10^. Pathological α-syn can also impair lysosomal degradation^10^, potentially creating a pathogenic cycle that promotes α-syn accumulation and neurodegeneration^11,12^. Thus, pathological α-syn may constitute both a cause and a consequence of autophagic dysfunction in PD^6^.

Biological sex has emerged as an important modifier of PD susceptibility and pathophysiology, with a higher prevalence in men^13^ and more pronounced pathological phenotypes in male animals in preclinical studies^14-17^. Potential contributors to the relative resilience observed in females may include the neuroprotective effects of estrogen^18-20^, and sex-related differences in mechanisms underlying PD pathogenesis, including inflammation and mitochondrial dysfunction^14,21^. Further characterization of cellular responses to α-syn-driven pathology across sexes will provide insight into the neurobiological basis of these differences.

We recently reported that male mice overexpressing A53T α-syn exhibit faster striatal axonopathy and earlier motor impairment than females^17^. Given the autophagy-lysosomal system’s central role in α-syn turnover, sex-dependent regulation of this pathway may contribute to differential vulnerability to α-syn pathology. Here, we investigated whether adeno-associated viral (AAV)-mediated A53T α-syn overexpression differentially alters the striatal autophagy-lysosomal system in male and female mice at 60 and 120 days after injection. To this end, we assessed autophagosome- and lysosome-associated structures by confocal immunofluorescence, quantified puncta area and density, and examined their association with α-syn expression and neurodegeneration.

## MATERIALS AND METHODS

### Animals and Experimental Design

This study used archived striatal tissue from a subset of mice included in our previous study^17^. 7-week-old C57BL/6J mice (14 males, 21-24 g; 15 females, 16-22 g; Janvier Labs) were housed in groups of 3-6 per individually ventilated cage under controlled conditions (22 ± 1°C, 55 ± 5% relative humidity, 12:12h light/dark cycle, lights on 08:00–20:00). Food and water were available *ad libitum*.

Sample sizes by sex, vector, and timepoint are provided in **Table S1**, and the experimental design is shown in **Figure 1A**. LC3 (microtubule-associated protein 1 light chain 3)/LAMP1 (lysosomal-associated membrane protein 1) immunofluorescence and image quantification were performed for this study. Previously published^17^ animal-level measurements of striatal α-syn and tyrosine hydroxylase (TH) immunoreactivity, together with axonal degeneration index (ADI) scores, were reanalyzed for the subset of animals included herein.

**Figure 1.**
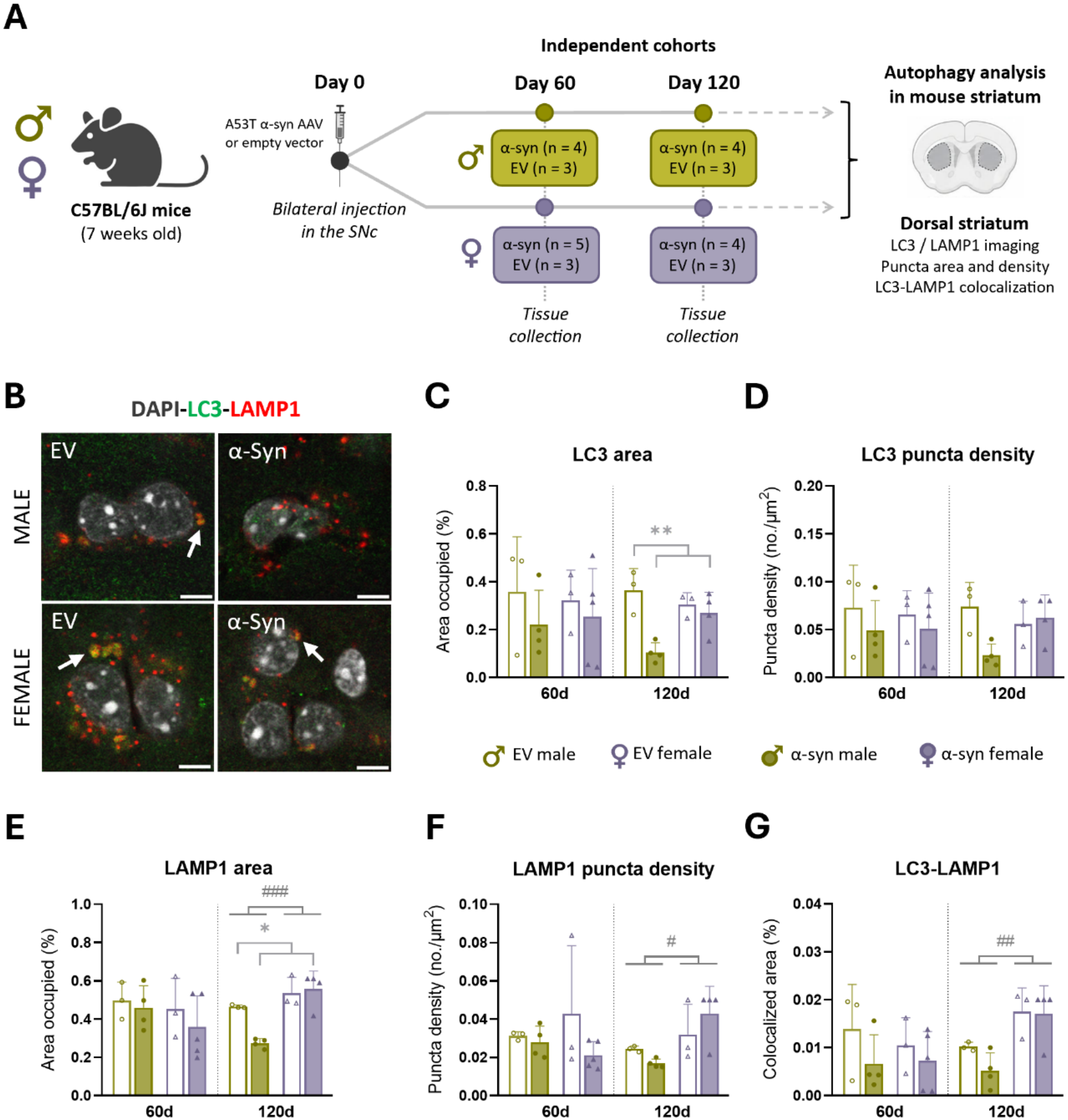
Experimental design and sex-dependent effect of alpha-synuclein overexpression on striatal autophagic-lysosomal markers. **(A)** Schematic representation of the experimental design and timepoints used for the striatal autophagy analysis. Adeno-associated viral (AAV) vectors encoding either mutant A53T human alpha-synuclein (α-syn) or an empty-AAV vector (EV) were bilaterally injected into the *substantia nigra pars compacta* (SNc) of male and female mice. **(B)** Representative confocal images of EV and α-syn mouse dorsal striatum at 120 days post-injection, showing immunofluorescent detection of the autophagosome marker LC3 (microtubule-associated protein 1 light chain 3, in green), the lysosomal marker LAMP1 (lysosomal-associated membrane protein 1, in red), and their combination (white arrows). Cell nuclei were visualized with DAPI (in white). Scale bar: 5 μm. **(C-G)** Quantification of autophagy-lysosomal markers in α-syn and EV mice at 60 and 120 days post-injection. **(C, E)** Total area occupied (%) by LC3 and LAMP1 puncta, respectively. **(D, F)** Density (puncta/µm^2^) of LC3 and LAMP1 puncta, respectively. **(G)** Percentage of LC3 puncta colocalized with LAMP1. Statistical significance was determined using two-way ANOVA, with significance levels represented by: * for the vector effect *p < 0.05; **p < 0.01; # the sex effect #p < 0.05; ##p < 0.01; ###p < 0.001.

### AAV Surgical Procedure

Stereotaxic injections were performed as previously described^17^. Briefly, recombinant AAV9 vectors (rAAV9-CMVie/SynP-WPRE) encoding either mutant A53T human α-syn (1 × 10^13^ genomic particles/mL) or an empty-AAV vector (3.1 × 10^12^ genomic particles/mL) were bilaterally injected into the *substantia nigra pars compacta* (1 μL per hemisphere). Coordinates relative to bregma were anteroposterior -2.9 mm; mediolateral ±1.4 mm; and dorsoventral -4.5 mm.

### Tissue Processing

Tissue collection and processing were performed as previously described^17^. At 60 or 120 days post-injection, animals were deeply anesthetized and transcardially perfused with saline followed by 4% paraformaldehyde in 0.1 M phosphate buffered saline (PBS). Brains were post-fixed (48h, 4°C), cryoprotected in 30% sucrose and 0.1% azide in 0.1 M PBS, coronally sectioned (40 μm), and stored at -20°C in cryoprotectant (30% ethylene glycol and 26% glycerol in 0.1 M PBS) until use.

### LC3 and LAMP1 Immunofluorescence

Free-floating striatal sections were blocked for 2h at room temperature in Tris-buffered saline (TBS) containing 3% bovine serum albumin (BSA; Sigma-Aldrich, A7906) and 0.1% Triton X-100, then incubated overnight at 4°C with rabbit anti-LC3 (1:500; Novus Biologicals, NB100-2220) and rat anti-LAMP1 (1:500; Developmental Studies Hybridoma Bank, 1D4B) antibodies diluted in TBS containing 3% BSA and 0.1% Tween-20. After washing, sections were incubated for 2h at room temperature with Alexa Fluor 488-conjugated goat anti-rabbit (1:500; Invitrogen, A32731TR) and Alexa Fluor 568-conjugated goat anti-rat (1:500; Invitrogen, A-11077) secondary antibodies, and 4’,6-diamidino-2-phenylindole (DAPI; 1:1000; Invitrogen, D1306) for nuclear counterstaining. Sections were mounted on Superfrost slides (Epredia™, 12302108) using Fluoromount-G.

### Neuropathological Measures

Previously acquired striatal α-syn and TH immunoreactivity measurements^17^, quantified as fiber optical density, were used to contextualize the autophagic-lysosomal findings.

Axonopathy severity in A53T α-syn mice was assessed using the previously described ADI, which combines striatal TH optical density and α-syn-positive axonal swelling counts^17^. Measures were standardized within each timepoint and combined to assign ordinal scores representing early (1), intermediate (2), or advanced (3) axonal degeneration. Original animal-level measurements and ADI scores were retained for the subset analyzed here.

### Confocal Imaging and Quantification of Autophagic-Lysosomal Markers

Images were acquired using a Leica STELLARIS 5 confocal microscope with a 63× oil-immersion objective (z-step, 0.34 μm) and constant acquisition settings. For each animal, 5-7 coronal sections spanning the rostrocaudal extent of the striatum were sampled, with one dorsal striatal field acquired per hemisphere per section. Mean z-stack depth was 7.52 μm per animal (n=29). TIFF images were analyzed in ImageJ (v2.9.0, v2.16.0) using custom macros to segment LC3- and LAMP1-positive puncta across complete z-stacks (sigma 0-10; thresholds: 35 (LC3) and 15 (LAMP1); minimum particle size, 0.032 µm^2^). Maximum particle sizes were 1 µm^2^ for LC3, and 5 µm^2^ for LAMP1 at 120 days, whereas no upper limit was applied for LAMP1 at 60 days. These limits excluded large structures identified as non-specific signals or imaging artifacts. Outcomes included the percentage of analyzed tissue area occupied by each marker and overlapping LC3/LAMP1 signals, excluding nuclear and vascular signals. Area measurements were obtained from complete z-stacks. For puncta-density measurements (puncta/μm^2^), optical planes were extracted at 15-plane intervals, fluorescence channels separated, and segmentation masks applied to quantify LC3- and LAMP1-positive puncta per unit area. Measurements across all sampled fields and sections from both hemispheres were averaged to obtain one value per animal for each outcome.

### Statistical Analysis

Statistical analyses were performed using R (v4.5.3) in RStudio (v2026.01.2). Continuous variables are summarized as mean (standard deviation, SD), with each animal constituting one independent experimental unit. Three-way ANOVAs examined the effects of vector, sex, time, and all interactions on autophagic-lysosomal markers. Separate two-way ANOVAs assessed vector, sex, and vector × sex effects at each timepoint for autophagic-lysosomal markers and neuropathological measures. Type III sums of squares with sum-to-zero contrasts were used to accommodate unequal sample sizes. Residual normality and homogeneity of variances were assessed using Shapiro-Wilk and Levene’s tests, respectively. Significant vector × sex interactions were followed by model-based comparisons using the pooled residual variance. Holm’s correction was applied jointly to four contrasts per outcome and timepoint.

Within the α-syn mice, separate linear models examined sex differences in autophagic-lysosomal markers adjusted for striatal α-syn optical density. Associations with striatal α-syn, TH, and the ADI were examined in A53T α-syn mice pooled across both timepoints using Spearman’s correlations and partial rank correlations adjusted for sex and time. Statistical significance was set at p < 0.05.

## RESULTS

### Sex-dependent alterations in striatal autophagic-lysosomal markers following alpha-synuclein overexpression

To characterize the organization of the striatal autophagy-lysosomal system, we quantified the area and density of LC3-positive (autophagosome-associated) and LAMP1-positive (lysosome-associated) puncta, together with their colocalization, providing a structural snapshot of these compartments and their spatial relationship.

We first examined the effects of vector, sex, and time on striatal autophagic-lysosomal markers using three-way ANOVA. Sex × time interactions were detected for LAMP1-positive area (F_1,21_ = 8.93, p = 0.007) and LC3-LAMP1 colocalization area (F_1,21_ = 7.35, p = 0.013), indicating that differences in autophagy-lysosomal parameters between timepoints varied by sex when averaged across experimental groups. A main effect of vector was also detected for LC3-positive area (F_1,21_ = 5.56, p = 0.028), with lower values in A53T α-syn mice when averaged across sexes and timepoints. No vector × sex × time interaction was detected for any marker **(Table S2)**.

We next examined the effects of α-syn overexpression at each timepoint using separate two-way ANOVAs including vector, sex, and their interaction. At 60 days, no significant vector, sex, or vector × sex effects were detected across the five measures **(Figure 1C-G; Table S3)**. At 120 days, α-syn overexpression was associated with lower LC3-positive area (F_1,10_ = 14.55, p = 0.003) and LAMP1-positive area (F_1,10_ = 5.76, p = 0.037) when averaged across sexes **(Figure 1B-G)**, corresponding to reductions of approximately 44% and 17% versus empty-AAV groups, respectively. However, these vector effects differed between sexes, as indicated by vector × sex interactions for LC3-positive area (F_1,10_ = 8.50, p = 0.015) and LAMP1-positive area (F_1,10_ = 9.22, p = 0.013). LC3 puncta density also showed a vector × sex interaction (F_1,10_ = 6.64, p = 0.028), although its main effect of vector was not significant (F_1,10_ = 3.70, p = 0.083).

Post-hoc comparisons showed that, relative to empty-AAV males, α-syn males exhibited approximately 71% lower LC3-positive area, 41% lower LAMP1-positive area, and 69% lower LC3 puncta density (Holm-adjusted p = 0.003, p = 0.010, and p = 0.039, respectively). The corresponding vector group comparisons in females were not significant **(Figure 1C-E; Table S4)**.

Main effects of sex were also detected at 120 days for LAMP1-positive area (F_1,10_ = 25.29, p < 0.001), LC3-LAMP1 colocalization area (F_1,10_ = 19.40, p = 0.001), and LAMP1 puncta density (F_1,10_ = 8.26, p = 0.017), with lower values in males when averaged across experimental groups **(Figure 1B-G; Table S3)**.

Together, these findings suggest sex- and time-dependent remodeling of striatal autophagosome- and lysosome-associated structures following α-syn overexpression, with reduced LC3/LAMP1 labeling most evident in males at 120 days.

### Relationship of autophagic-lysosomal alterations with alpha-synuclein expression and axonopathy severity

To provide pathological context for the autophagic-lysosomal findings, we reanalyzed previously published striatal α-syn and TH measurements^17^ from the animals included in this study. At 60 days, α-syn immunoreactivity was higher in α-syn mice than in the empty-AAV controls (F_1,11_ = 27.7, p < 0.001), whereas TH immunoreactivity showed no vector effect (F_1,11_ = 0.08, p = 0.779). At 120 days, α-syn immunoreactivity remained elevated (vector effect: F_1,10_ = 28.5, p < 0.001; **Figure 2A)**, and TH immunoreactivity was lower in α-syn mice (F_1,10_ = 5.11, p = 0.047; **Figure 2B**). Neither marker showed a significant vector × sex interaction at either timepoint **(Table S3)**.

**Figure 2.**
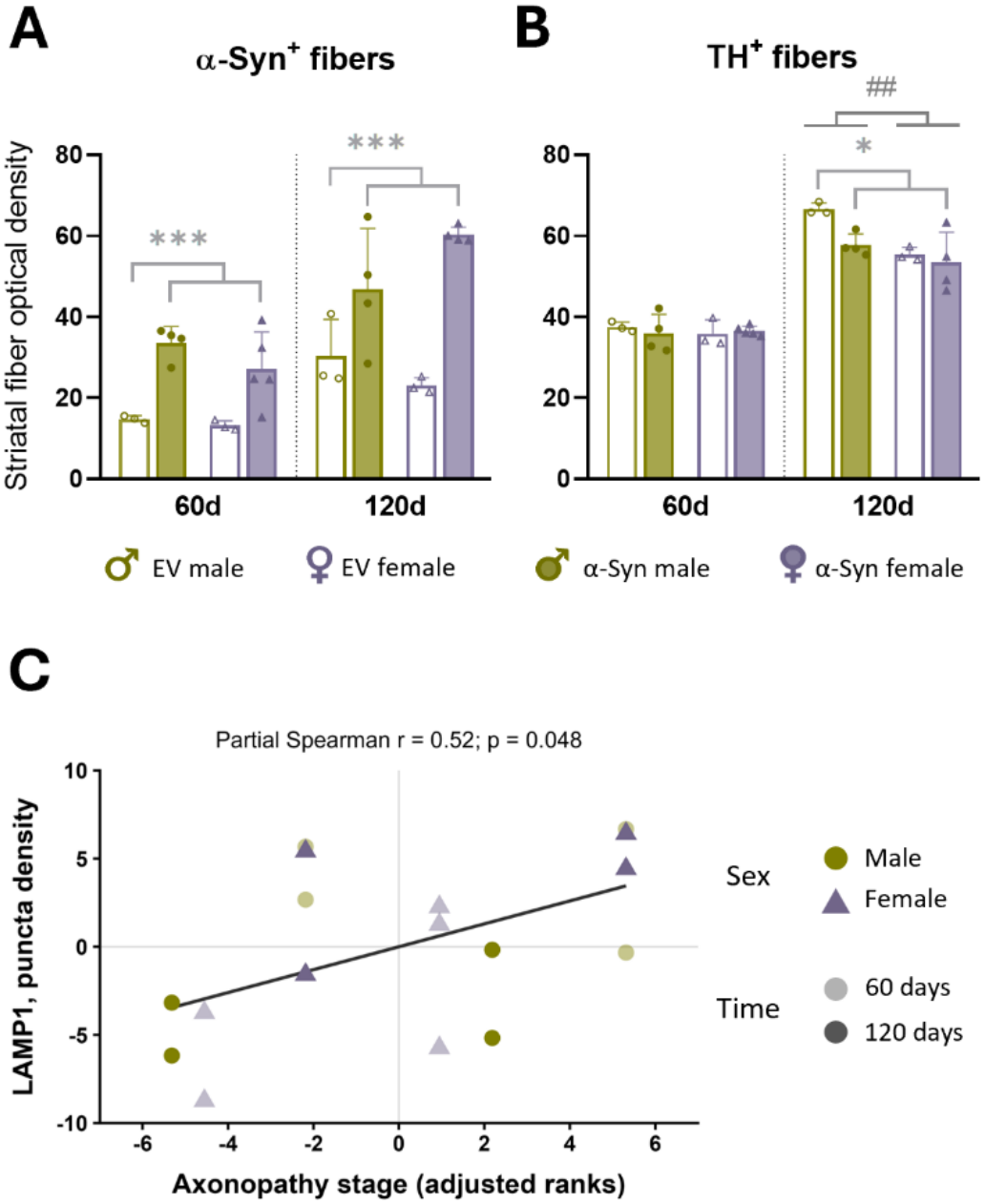
Pathological context and relationship to axonopathy. Quantification of the optical density of alpha-synuclein-positive (α-syn^+^) fibers **(A)** and dopaminergic fibers **(B)** in the striatum in male and female A53T α-syn and empty-AAV vector (EV) mice at 60 and 120 days post-injection. Dopaminergic neurons were positive for tyrosine hydroxylase (TH^+^). Statistical significance was determined using two-way ANOVA, with significance levels represented by: * for the vector effect *p < 0.05; ***p < 0.001; # the sex effect ##p < 0.01. **(C)** Partial Spearman correlations between axonopathy stage and LAMP1 puncta density in α-syn animals pooled across both timepoints (n = 17), adjusting for sex and time. Each point represents one animal. Axes display residuals obtained by regressing the ranks of each variable on sex and time; negative values therefore indicate ranks below those predicted by the covariates. Black line shows linear fit to the residualized ranks across all animals.

To assess whether sex differences in autophagic-lysosomal markers persisted after accounting for variation in α-syn immunoreactivity, we fitted exploratory linear models within the α-syn group at 120 days. After adjustment, males showed lower LC3-positive area (β_male_ [standard error, SE] = -0.17 [0.07], p = 0.046), LAMP1-positive area (β_male_ [SE] = -0.30 [0.07], p = 0.006), LC3-LAMP1 colocalization area (β_male_ [SE] = -0.014 [0.004], p = 0.024), and LAMP1 puncta density (β_male_ [SE] = -0.026 [0.010], p = 0.047) than females. The adjusted difference in LC3 puncta density was not significant (p = 0.069; **Table S5**).

Finally, we examined associations between autophagic-lysosomal markers and striatal pathology within the α-syn group. The higher TH signal observed at 120 days could partly reflect increased axonal swelling despite ongoing pathology. As TH immunoreactivity alone may not fully capture axonal degeneration, we used the ADI^17^, which combines striatal TH optical density with α-syn-positive axonal swelling counts, thereby assessing complementary features of axonal degeneration. Across both timepoints (n=17), greater LAMP1 puncta density was associated with a higher ADI after adjustment for sex and time (partial rank correlation = 0.518, p = 0.048; **Figure 2C**). No other partial correlation reached significance, including associations with striatal α-syn or TH immunoreactivity **(Table S6)**.

## DISCUSSION

This study identifies sex-dependent alterations in striatal autophagic-lysosomal markers in an A53T α-syn model of PD. At 120 days, α-syn males exhibited reduced LC3- and LAMP1-positive areas and lower LC3 puncta density relative to male controls. Several differences between α-syn males and females persisted after accounting for striatal α-syn immunoreactivity. Furthermore, greater LAMP1 puncta density was associated with more advanced axonopathy after adjustment for sex and time, linking lysosomal alterations to pathological severity. Together, these observations extend our previous finding of faster axonal degeneration in males^17^ by showing that biological sex influences the striatal autophagic-lysosomal phenotype accompanying α-syn pathology.

Autophagic-lysosomal alterations have been documented in both experimental and human PD. Dehay et al. demonstrated lysosomal depletion preceding autophagosome accumulation and dopaminergic degeneration in MPTP models, alongside reduced nigral LAMP1 levels in *postmortem* PD tissue^9^. In an AAV α-syn rat model, Decressac et al. identified impaired nuclear translocation of transcription factor EB (TFEB), a regulator of lysosomal biogenesis, and reported a similar alteration in human PD midbrain^22^. Human *postmortem* studies have also shown reductions in LAMP2A and HSC70, key components of chaperone-mediated autophagy^23^. Our findings extend this evidence to the dorsal striatum, where reductions in LC3 and LAMP1 labeling following α-syn overexpression were detected in males at 120 days. The accompanying vector × sex interactions indicate that sex modifies this response, adding a biological source of variation to the characterization of autophagic-lysosomal alterations in α-syn models.

The persistence of sex differences after adjustment for α-syn immunoreactivity suggests that the observed phenotype is not explained solely by differences in expression levels. This raises the possibility of sex-dependent regulation of autophagic-lysosomal compartments during sustained α-syn overexpression. Supporting a role for sex-related mechanisms in α-syn homeostasis, Rajsombath et al. reported delayed motor abnormalities in female 3K α-syn transgenic mice and beneficial effects of brain-selective estrogen treatment on α-syn tetramerization and the motor phenotype^24^. Together, these observations provide a rationale for investigating sex-dependent regulation of protein handling as a potential contributor to differential responses to α-syn pathology.

The association with axonopathy further connects the striatal lysosomal phenotype to tissue injury. Among α-syn animals, greater LAMP1 puncta density accompanied more advanced axonal pathology, suggesting that lysosomal compartments undergo remodeling as axonal damage increases. This relationship addresses variation in pathological severity within α-syn animals, whereas the lower LAMP1-positive area in males describes a difference between experimental groups. The findings highlight complementary aspects of the phenotype and support assessing lysosomal organization alongside axonal integrity when evaluating the response to α-syn. This combined perspective is relevant to therapies targeting autophagic-lysosomal pathways. In α-syn models, TFEB overexpression and pharmacological enhancement of autophagy promoted α-syn clearance and protected dopaminergic neurons^22^. Our findings add a rationale for determining whether these interventions produce comparable cellular and neuroprotective effects across sexes. Assessing lysosomal function together with axonal integrity in both sexes would help establish whether modifying this pathway alters the course of axonal injury and whether treatment responses depend on the sex-related differences identified here.

Limitations include small, unbalanced subgroups, cross-sectional sampling, restricted dorsal striatal coverage, and the exploratory nature of the adjusted models and correlation analyses. Moreover, static LC3/LAMP1 measurements do not establish autophagic flux or degradative efficiency, and the absence of cell-type-specific labeling prevents distinguishing neuronal from glial contributions. Strengths include the incorporation of both sexes and the complementary assessment of autophagy-lysosomal markers, α-syn expression, and axonopathy severity within a single experimental framework. The observed sex-related differences in autophagy-lysosomal marker profiles warrant consideration when developing therapies to preserve nigrostriatal axons.

## STATEMENTS

### Data availability

Data supporting this study are available on the institutional open-access repository of the University of the Basque Country (EHU) ADDI, at: https://doi.org/10.82518/M4KZZW. For further inquiries, please contact the corresponding author.

### Ethical Approval

All experimental procedures were approved by the Local Ethical Committee for Animal Research of the University of the Basque Country (EHU, CEEA, protocols M20/2019/219, M20-2022-215, and M30/2019/220) and were conducted in accordance with the European Directive 2010/63/EU and Spanish Royal Decree 53/2013 regulating the use of laboratory animals.

### Author Contributions

R.B. contributed to Investigation, Data curation, Formal analysis, Visualization, and Writing – review & editing; M.Z. contributed to Investigation, Methodology, Data curation, Formal analysis, Visualization, and Writing – review & editing; E.E. contributed to Investigation, Data curation, and Formal analysis; I.J.-R. contributed to Investigation and Visualization; A.M.-G. contributed to Data curation, Formal analysis, and Writing – review & editing; and J.-A.R.O. contributed to Investigation and Methodology; J.V.L. contributed to Funding acquisition and Writing – review & editing; N.P.M. contributed to Conceptualization, Supervision, and Writing – review & editing; A.P.-Z. contributed to Conceptualization, Methodology, Project administration, Supervision, Funding acquisition, Writing – original draft, and Writing – review & editing; T.M.-H. contributed to Conceptualization, Methodology, Project administration, Supervision, Writing – original draft, and Writing – review & editing. All authors contributed to data interpretation, critically revised the manuscript, and approved the final version.

## Acknowledgements

The authors thank Dr. Benjamin Dehay (Univ. Bordeaux, CNRS, IMN, UMR 5293, F-33000 Bordeaux, France) for providing the viral vectors used in this study, and the Achucarro Basque Center for Neuroscience-Imaging Facility (EHU Scientific Park, Spain) for access to the Leica STELLARIS 5 confocal microscope.

