## Supplementary material for "Alpha-synuclein overexpression induces sex-dependent alterations in striatal autophagic-lysosomal markers in a Parkinson’s disease mouse model": Table S1; Table S2; Table S3; Table S4; Table S5; Table S6

[LA]A. Plaza-Zabala and T. Morera-Herreras should be considered shared last authors.

**Table of contents:**

**Table S1.** Sample sizes and descriptive statistics for striatal autophagic-lysosomal and pathological markers

**Table S2.** Three-way ANOVA of striatal autophagic-lysosomal markers

**Table S3.** Two-way ANOVA of striatal autophagic-lysosomal and pathological markers at 60 and 120 days

**Table S4.** Post-hoc comparisons following significant vector × sex interactions at 120 days

**Table S5.** Sex differences in striatal autophagic-lysosomal markers adjusted for alpha-synuclein immunoreactivity at 120 days

**Table S6**. Associations between striatal autophagic-lysosomal markers and pathological measures in A53T alpha-synuclein mice

**Table S1. Sample sizes and descriptive statistics for striatal autophagic-lysosomal and pathological markers**

| | Empty-AAV, female | Empty-AAV, male | A53T α-syn, female | A53T α-syn, male |
|---|---|---|---|---|
| **60 days post-injection** | | | | |
| Animals, n | 3 | 3 | 5 | 4 |
| LC3-positive area (%) | 0.323 (0.126) | 0.358 (0.230) | 0.253 (0.201) | 0.221 (0.143) |
| LC3 puncta density (puncta/μm²) | 0.066 (0.025) | 0.073 (0.045) | 0.051 (0.037) | 0.049 (0.031) |
| LAMP1-positive area (%) | 0.454 (0.159) | 0.496 (0.096) | 0.358 (0.162) | 0.459 (0.115) |
| LAMP1 puncta density (puncta/μm²) | 0.043 (0.036) | 0.031 (0.002) | 0.021 (0.007) | 0.028 (0.008) |
| LC3-LAMP1 colocalization area (%) | 0.010 (0.006) | 0.014 (0.009) | 0.007 (0.006) | 0.007 (0.006) |
| Striatal TH optical density | 35.80 (3.44) | 37.51 (1.20) | 36.47 (1.21) | 35.92 (4.73) |
| Striatal α-syn optical density | 13.15 (1.21) | 14.75 (0.86) | 27.19 (9.06) | 33.53 (4.10) |
| ADI, early/mild/advanced, n | N/A | N/A | 2 / 3 / 0 | 0 / 2 / 2 |
| **120 days post-injection** | | | | |
| Animals, n | 3 | 3 | 4 | 4 |
| LC3-positive area (%) | 0.305 (0.050) | 0.363 (0.091) | 0.271 (0.088) | 0.105 (0.043) |
| LC3 puncta density (puncta/μm²) | 0.055 (0.023) | 0.074 (0.025) | 0.063 (0.024) | 0.023 (0.011) |
| LAMP1-positive area (%) | 0.536 (0.083) | 0.465 (0.009) | 0.559 (0.095) | 0.273 (0.025) |
| LAMP1 puncta density (puncta/μm²) | 0.032 (0.017) | 0.024 (0.002) | 0.043 (0.014) | 0.017 (0.002) |
| LC3-LAMP1 colocalization area (%) | 0.018 (0.005) | 0.010 (0.001) | 0.017 (0.006) | 0.005 (0.003) |
| Striatal TH optical density | 55.39 (1.73) | 66.65 (1.48) | 53.43 (7.45) | 57.71 (2.72) |
| Striatal α-syn optical density | 23.03 (1.94) | 30.35 (9.02) | 60.24 (1.92) | 46.75 (15.07) |
| ADI, early/mild/advanced, n | N/A | N/A | 0 / 2 / 2 | 0 / 2 / 2 |

Values are presented as mean (SD), except for sample sizes and axonal degeneration index, which are reported as counts. The analysis cohort comprised 29 animals with autophagic-lysosomal marker measurements: 15 mice at 60 days and 14 mice at 120 days post-injection. All continuous outcomes were available for every animal; therefore, the sample sizes shown in the first row apply to all continuous outcomes within each panel. Each observation represents one animal. The ADI is reported as the number of animals assigned to early, mild or advanced stage, respectively, and was available for all 17 A53T α-syn animals. Striatal TH and α-syn measurements were reused from our previous study (Zubelzu et al., 2026, *Cell Mol Neurobiol*, 10.1007/s10571-026-01707-9); the descriptive statistics presented here were recalculated for the subset of animals included in the present study. α-Syn, alpha-synuclein; ADI, Axonal Degeneration Index; LAMP1, lysosomal-associated membrane protein 1; LC3, microtubule-associated protein 1 light chain 3; N/A, not applicable; SD, standard deviation; TH, tyrosine hydroxylase.

**Table S2. Three-way ANOVA of striatal autophagic-lysosomal markers**

| Outcome | Model term | F | p-value |
|---|---|---|---|
| LC3-positive area | Vector | 5.562 | **0.028** |
| | Sex | 0.240 | 0.629 |
| | Time | 0.276 | 0.605 |
| | Vector × sex | 1.902 | 0.182 |
| | Vector × time | 0.169 | 0.685 |
| | Sex × time | 0.277 | 0.604 |
| | Vector × sex × time | 0.545 | 0.469 |
| LC3 puncta density | Vector | 3.422 | 0.078 |
| | Sex | 0.121 | 0.732 |
| | Time | 0.271 | 0.608 |
| | Vector × sex | 2.297 | 0.145 |
| | Vector × time | 0.014 | 0.906 |
| | Sex × time | 0.354 | 0.558 |
| | Vector × sex × time | 1.259 | 0.275 |
| LAMP1-positive area | Vector | 3.268 | 0.085 |
| | Sex | 1.611 | 0.218 |
| | Time | 0.151 | 0.701 |
| | Vector × sex | 0.878 | 0.359 |
| | Vector × time | 0.050 | 0.825 |
| | Sex × time | 8.926 | **0.007** |
| | Vector × sex × time | 2.669 | 0.117 |
| LAMP1 puncta density | Vector | 1.053 | 0.316 |
| | Sex | 3.339 | 0.082 |
| | Time | 0.087 | 0.770 |
| | Vector × sex | <0.001 | 0.998 |
| | Vector × time | 1.731 | 0.202 |
| | Sex × time | 1.930 | 0.179 |
| | Vector × sex × time | 2.908 | 0.103 |
| LC3-LAMP1 colocalization area | Vector | 3.782 | 0.065 |
| | Sex | 4.200 | 0.053 |
| | Time | 1.869 | 0.186 |
| | Vector × sex | 1.082 | 0.310 |
| | Vector × time | 0.217 | 0.646 |
| | Sex × time | 7.351 | **0.013** |
| | Vector × sex × time | 0.013 | 0.909 |

Exploratory three-way ANOVA results for vector (empty-AAV or A53T α-syn), sex (female or male), time (60 or 120 days), and all interactions. All factors were treated as categorical variables, and analyses used Type III sums of squares with sum-to-zero contrasts. Each animal contributed one observation per outcome. α-Syn, alpha-synuclein; LAMP1, lysosomal-associated membrane protein 1; LC3, microtubule-associated protein 1 light chain 3.

**Table S3. Two-way ANOVA of striatal autophagic-lysosomal and pathological markers at 60 and 120 days**

| | Day | Vector | | Sex | | Interaction | |
|---|---|---|---|---|---|---|---|
| | | F | p-value | F | p-value | F | p-value |
| ***Autophagic-lysosomal markers*** | | | | | | | |
| LC3-positive area | 60 | 1.16 | 0.305 | <0.001 | 0.984 | 0.13 | 0.730 |
| | 120 | 14.5 | **0.003** | 1.96 | 0.192 | 8.50 | **0.015** |
| LC3 puncta density | 60 | 1.06 | 0.325 | 0.02 | 0.886 | 0.05 | 0.819 |
| | 120 | 3.70 | 0.083 | 0.85 | 0.379 | 6.64 | **0.028** |
| LAMP1-positive area | 60 | 0.81 | 0.389 | 0.95 | 0.351 | 0.16 | 0.700 |
| | 120 | 5.76 | **0.037** | 25.29 | **<0.001** | 9.22 | **0.013** |
| LAMP1 puncta density | 60 | 2.07 | 0.178 | 0.07 | 0.793 | 1.10 | 0.316 |
| | 120 | 0.07 | 0.801 | 8.26 | **0.017** | 2.31 | 0.159 |
| LC3-LAMP1 colocalization area | 60 | 2.14 | 0.172 | 0.16 | 0.696 | 0.31 | 0.586 |
| | 120 | 1.87 | 0.201 | 19.40 | **0.001** | 1.14 | 0.310 |
| ***Pathological markers*** | | | | | | | |
| Striatal TH optical density | 60 | 0.08 | 0.779 | 0.13 | 0.724 | 0.51 | 0.490 |
| | 120 | 5.11 | **0.047** | 10.39 | **0.009** | 2.10 | 0.178 |
| Striatal α-syn optical density | 60 | 27.7 | **<0.001** | 1.63 | 0.229 | 0.58 | 0.463 |
| | 120 | 28.5 | **<0.001** | 0.38 | 0.552 | 4.30 | 0.065 |

Two-way ANOVA results for vector, sex, and their interaction at each time point. All factors were treated as categorical variables, and analyses used Type III sums of squares with sum-to-zero contrasts. Each animal contributed one observation per outcome. α-Syn, alpha-synuclein; LAMP1, lysosomal-associated membrane protein 1; LC3, microtubule-associated protein 1 light chain 3; TH, tyrosine hydroxylase.

**Table S4. Post-hoc comparisons following significant vector × sex interactions at 120 days**

| Outcome | Contrast | Estimated difference (95% CI) | Holm-adjusted p-value |
|---|---|---|---|
| LC3-positive area | A53T α-syn – empty-AAV, females | −0.03 (−0.16 to 0.09) | 0.679 |
| | A53T α-syn – empty-AAV, males | −0.26 (−0.38 to −0.14) | **0.003** |
| | Male − female, empty-AAV | 0.06 (−0.07 to 0.19) | 0.679 |
| | Male − female, A53T α-syn | −0.17 (−0.28 to −0.05) | **0.024** |
| LAMP1-positive area | A53T α-syn – empty-AAV, females | 0.02 (−0.09 to 0.13) | 0.662 |
| | A53T α-syn – empty-AAV, males | −0.19 (−0.30 to −0.08) | **0.010** |
| | Male − female, empty-AAV | −0.07 (−0.19 to 0.05) | 0.434 |
| | Male − female, A53T α-syn | −0.29 (−0.39 to −0.18) | **<0.001** |
| LC3 puncta density | A53T α-syn – empty-AAV, females | 0.007 (−0.03 to 0.04) | 0.654 |
| | A53T α-syn – empty-AAV, males | −0.05 (−0.09 to −0.02) | **0.039** |
| | Male − female, empty-AAV | 0.02 (−0.020 to 0.06) | 0.598 |
| | Male − female, A53T α-syn | −0.04 (−0.073 to −0.01) | 0.070 |

Model-based contrasts for outcomes with a significant vector × sex interaction at 120 days. Differences are expressed as the first condition minus the second condition listed and were estimated using the pooled residual variance from the corresponding two-way model (residual df = 10). For each outcome, Holm adjustment was applied jointly to four contrasts: the vector effect within each sex and the sex difference within each vector group. Confidence intervals are unadjusted 95% intervals. Area differences are expressed in percentage points, and puncta-density differences in puncta/$\mu m^2$. Bold p-values indicate $p < 0.05$. α-Syn, alpha-synuclein; CI, confidence interval; LAMP1, lysosomal-associated membrane protein 1; LC3, microtubule-associated protein 1 light chain 3.

**Table S5. Sex differences in striatal autophagic-lysosomal markers adjusted for alpha-synuclein immunoreactivity at 120 days**

| Outcome | β | SE | 95% CI | p-value |
|---|---|---|---|---|
| LC3-positive area | −0.175 | 0.066 | −0.344 to −0.005 | **0.046** |
| LC3 puncta density | −0.042 | 0.018 | −0.088 to 0.005 | 0.069 |
| LAMP1-positive area | −0.302 | 0.065 | −0.470 to −0.134 | **0.006** |
| LAMP1 puncta density | −0.026 | 0.010 | −0.051 to −0.0004 | **0.047** |
| LC3-LAMP1 colocalization area | −0.014 | 0.004 | −0.025 to −0.003 | **0.024** |

Analyses were restricted to A53T α-syn animals at 120 days post-injection (4 females and 4 males). Separate ordinary least-squares linear models were fitted for each outcome, with sex and striatal α-syn optical density as predictors. β represents adjusted differences as males minus females; negative values indicate lower values in males after adjustment for α-syn immunoreactivity. Area differences are expressed in percentage points, and puncta-density differences in puncta/$\mu m^2$. Standard errors, two-sided p-values, and 95% confidence intervals were calculated using the conventional model-based variance estimator and five residual degrees of freedom. p-values and confidence intervals were not adjusted for multiple testing across the five outcomes. These analyses were exploratory. Significant p-values (<0.05) are highlighted in bold. α-Syn, alpha-synuclein; CI, confidence interval; LAMP1, lysosomal-associated membrane protein 1; LC3, microtubule-associated protein 1 light chain 3; SE, standard error.

**Table S6. Associations between striatal autophagic-lysosomal markers and pathological measures in A53T alpha-synuclein mice**

| Autophagic-lysosomal outcome | Pathological measure | ρ | Unadjusted p-value | Partial rank correlation | Partial correlation p-value |
|---|---|---|---|---|---|
| LC3-positive area | TH optical density | −0.127 | 0.626 | −0.023 | 0.936 |
| | α-Syn optical density | 0.216 | 0.406 | 0.471 | 0.076 |
| | ADI | −0.022 | 0.934 | 0.140 | 0.619 |
| LC3 puncta density | TH optical density | −0.108 | 0.680 | −0.027 | 0.923 |
| | α-Syn optical density | 0.240 | 0.353 | 0.478 | 0.072 |
| | ADI | −0.001 | 0.996 | 0.143 | 0.612 |
| LAMP1-positive area | TH optical density | −0.112 | 0.670 | −0.382 | 0.161 |
| | α-Syn optical density | 0.338 | 0.184 | 0.408 | 0.131 |
| | ADI | 0.267 | 0.300 | 0.385 | 0.156 |
| LAMP1 puncta density | TH optical density | −0.137 | 0.599 | −0.391 | 0.150 |
| | α-Syn optical density | 0.328 | 0.198 | 0.420 | 0.119 |
| | ADI | 0.369 | 0.145 | 0.518 | **0.048** |
| LC3-LAMP1 colocalization area | TH optical density | 0.156 | 0.551 | −0.174 | 0.536 |
| | α-Syn optical density | 0.450 | 0.070 | 0.393 | 0.147 |
| | ADI | 0.146 | 0.576 | 0.242 | 0.385 |

Analyses included 17 A53T α-syn animals pooled across both time points: 9 mice at 60 days (5 females and 4 males) and 8 mice at 120 days (4 females and 4 males). All variables were available for every animal. Unadjusted associations were assessed using Spearman's rank correlation. Partial rank correlations were calculated by assigning average ranks to tied observations, regressing the ranks of each variable on sex and time as categorical covariates, and calculating the Pearson correlation between the resulting residuals. Two-sided p-values were obtained using a t approximation with 15 degrees of freedom for unadjusted correlations and 13 degrees of freedom for partial correlations. Bold indicates $p < 0.05$. α-Syn, alpha-synuclein; ADI, Axonal Degeneration Index; LAMP1, lysosomal-associated membrane protein 1; LC3, microtubule-associated protein 1 light chain 3; N/A, not applicable; SD, standard deviation; TH, tyrosine hydroxylase.
